# Symmetry breaking and avalanche shapes in modular neural networks

**DOI:** 10.1101/2025.10.29.685308

**Authors:** Antonio de Candia, Davide Conte, Hanieh Alvankar Golpayegan, Silvia Scarpetta

## Abstract

Modularity is as a key characteristic of structural and functional brain networks across species and spatial scales. We investigate the stochastic Wilson-Cowan model on a modular network in which synaptic strengths differ between intra-module and inter-module connections. The system exhibits a rich phase diagram comprising symmetric (SL and SH) and “broken symmetry” (B1, B2, …) phases. The phase SL (SH) is characterized by the same low (high) activity in all the modules, while the B*m* phases are characterized by a high activity in *m* modules and low activity in the remaining modules. Between SL and SH, and between SL and B1, there are two lines of critical points, where the system shows a critical behaviour, with power law distributions in the avalanches. Along these lines, avalanche shapes differ systematically: they are symmetric or right-skewed at the SL-SH transition, but can become left-skewed over intermediate durations along the SL-B1 critical line. These results provide a theoretical framework that accounts for both symmetric and left-skewed neural avalanche shapes observed experimentally, linking modular organization to critical brain dynamics.

## I. INTRODUCTION

Spontaneous brain activity unfolds over multiple spatial and temporal scales, giving rise to rich dynamical patterns that include intermittent bursts known as neuronal avalanches [1–6]. First identified in rat cortical slices, and later confirmed in vivo across rodents [7, 8], non-human primates [9–12] and humans [13–19], these cascades display size and lifetime distributions devoid of a characteristic scale, a signature long interpreted as evidence that cortical networks reside near a nonequilibrium critical point [20]. Scale-free avalanches were reproduced also in many computational models of neural activity [21–27], among which in a stochastic version of the Wilson-Cowan model [28–33].

These scaling laws place the brain within the broader realm of crackling noise, in which slowly driven systems respond through impulsive events spanning many orders of magnitude [34, 35], ranging from earthquakes [36, 37] to Barkhausen noise in ferromagnets [38–40]. Historically, the power-law distributions of avalanche sizes and durations have been considered hallmarks of criticality, however this could not be a sufficient indicator of criticality, as they can arise from various non-critical mechanisms [41, 42].

To overcome the ambiguity of power-law exponents alone, recent research has increasingly focused on the mean temporal profiles (shapes) of neuronal avalanches as a more stringent and reliable test for criticality [43–45]. Critical systems predict that when appropriately rescaled, the mean temporal profiles of avalanches of widely varying durations should collapse onto a single universal scaling function, often approximated by an inverted parabola [34].

Experimental observations, however, have yielded mixed results regarding these shapes. A parabolic profile for avalanches in the scaling regime has been observed with two-photon imaging of neurons in the superficial cortex of awake mice [46, 47]. Nearly symmetric avalanche shapes have been observed in intracranial depth recordings in humans [48], as well as in the spiking activity of the mammalian primary visual cortex, both in anesthetized animals (rats and monkeys) and freely moving mice, where leftward asymmetric avalanches tend to occur at lower values of the spiking variability [49]. In nonhuman primates, chronically implanted high-density microelectrode arrays showed a nearly parabolic avalanche shape, modulated by *γ*–scillations at intermediate raster plot binning [12, 50].

Many other experimental measurements have revealed clear departures from perfect symmetry, displaying leftward skewing and extended tails. Whole-brain light-sheet imaging of transgenic zebrafish larvae has shown that average avalanche profiles collapse into a distinctly leftward-asymmetric form, characterized by rapid initiation and slow decay [51]. Similar leftward asymmetry has been observed in high-resolution in vitro recordings of dissociated cortical cultures [52], whereas organotypic cortical cultures exhibit collapse onto approximately inverted parabolic profiles with only minimal leftward skewing. This characteristic leftward asymmetry has also been reported in human EEG recordings, both during recovery from hypoxia [53] and in preterm infants [54]. Remarkably, EEG data from preterm infants revealed systematic changes in avalanche shape asymmetry as a function of avalanche duration, with avalanches transitioning from symmetric to leftward-asymmetric forms. Moreover, the degree of asymmetry at intermediate durations correlated significantly with long-term neurodevelopmental outcomes [54]. Similar asymmetric profiles are also observed in physical crackling noise systems, such as Barkhausen noise [55] and earthquakes [56, 57], where they have been linked to complex underlying mechanisms like “effective mass” or transient forces counteracting propagation [58]. The origins of this leftward asymmetry in biological data remain puzzling. A notable contrast arises from the simple (non-modular) stochastic Wilson-Cowan model, where symmetric or even right-skewed avalanche profiles have been found [29, 30].

Beyond the intrinsic dynamics of neuronal populations, the structural organization of brain networks plays a fundamental role in shaping neuronal activity. The brain is widely recognized as a complex network characterized by a modular and often hierarchically modular architecture [59–62]. Modules are typically defined as subsets of nodes, often corresponding to anatomically or functionally related neuronal regions, that are densely and strongly interconnected internally, while maintaining comparatively sparser and weaker connections with nodes in other modules [63]. The role of modular network topology in spontaneous collective dynamics and the emergence of critical behavior has been investigated in previous studies [64–73], but the impact of modularity on avalanche shape and on the manner in which avalanches propagate across modular regions remains an important open question. Motivated by these unresolved questions and the persistent discrepancies between current models and experimental findings regarding avalanche shapes, our study aims to bridge this gap by investigating neuronal avalanche distributions and, critically, their mean temporal shapes within a stochastic Wilson-Cowan model incorporating a modular network topology.

We first show that, on a modular network, the Wilson-Cowan model shows different phases, depending on the strength of excitatory and inhibitory connections between modules and within each module. Phases can be symmetric (SL and SH) characterized by the same (high and low) mean activity on all the modules, or they can be characterized by a breaking of the symmetry between modules (B*m*, with *m* = 1, 2,…) corresponding to *m* modules having a high activity, and the others a low activity. Between SL and SH, and between SL and B1, there are two lines of critical points, where the system displays a scale-free distribution of the avalanches.

## II. WILSON-COWAN MODEL ON A MODULAR NETWORK

We consider the stochastic Wilson-Cowan model [28]. In this model, neurons have two possible states, namely active and quiescent, and make transitions from one state to the other with specified rates. The transition from active to quiescent takes place with a constant rate *α*, while that from quiescent to active with a rate *f* (*s*_*i*_), where *s*_*i*_ is the input of the *i*-th neuron that we are considering, and *f* (*s*) is an activation function taken equal to

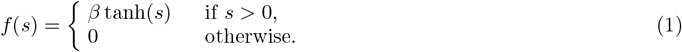

The input of the *i*-th neuron is given by

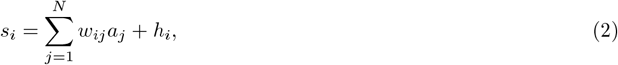

where *a*_*j*_ = 0, 1 if the *j*-th neuron is quiescent or active, *w*_*ij*_ is the strength of the connection going from *j* to *i*, and *h*_*i*_ is the external input to neuron *i*. In the simplest case, one takes *N*_*E*_ excitatory and *N*_*I*_ inhibitory neurons, with *N*_*E*_ + *N*_*I*_ = *N*, and the connections *w*_*ij*_ having only two values, namely *w*_*E*_*/N*_*E*_ if the pre-synaptic neuron *j* is excitatory, or −*w*_*I*_*/N*_*I*_ if *j* is inhibitory.

It was shown in [29] that, when the external input *h*_*i*_ goes to zero, the dynamics of the model undergoes a transition (bifurcation) at a critical value of the imbalance *w*_0_ = *w*_*E*_ −*w*_*I*_, namely *w*_0*c*_ = *β*^−1^*α*. For *w*_0_ *< w*_0*c*_, the fraction of active neurons goes to zero when *h*_*i*_ → 0, while for *w*_0_ *> w*_0*c*_ it is finite also for *h*_*i*_ → 0, so that there is a self-sustained activity. Exactly at *w*_0_ = *w*_0*c*_, fluctuations are maximized, and the dynamics is characterized by bursts of activity (avalanches), whose distribution is scale-free, with the critical exponents of the branching model. The transition can be understood as a “transcritical bifurcation”, where the quiescent fixed point of the dynamics becomes unstable, and the self-sustained fixed point becomes stable. As such, it can be found looking when one of the eigenvalues of the stability matrix becomes equal to zero.

In this paper, we consider a version of the model in which the network is composed by *M* modules, each containing *N*_*E*_ excitatory and *N*_*I*_ inhibitory neurons. Each neuron in module *k* receives input from excitatory neurons of module *l* with synaptic strengths 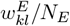, and from inhibitory neurons of module *l* with synaptic strengths 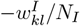, so that the input *s*_*k*_ of neurons in module *k* is

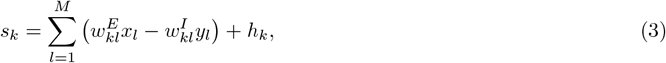

where *x*_*l*_ and *y*_*l*_ are the fractions of excitatory and inhibitory active neurons in module *l* (activities), *h*_*k*_ is the external input to module *k*. In the Gaussian noise approximation [28, 74], the dynamics of the network is determined by the equations

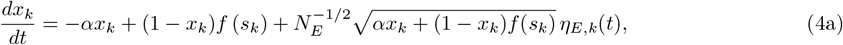

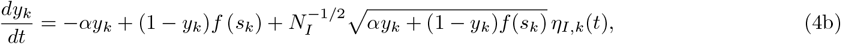

where *η*_*E,k*_(*t*), *η*_*I,k*_(*t*) are normal, Gaussian independent white noises. The variables *x*_*k*_ and *y*_*k*_ can be written as 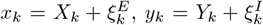, where *X*_*k*_ and *Y*_*k*_ are the “deterministic” components of the activities, that obey the deterministic equations

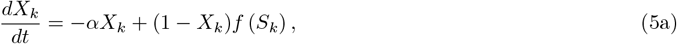

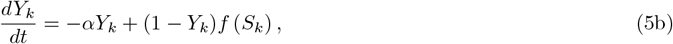

with 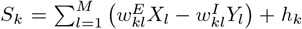. The deterministic components evolve until they reach a stable (attractive) fixed point, that is a configuration of *X*_*k*_ and *Y*_*k*_ that set to zero the right hand sides of Eqs. (5). We can then define 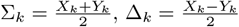, where Σ_*k*_ represents the total activity of the *k*-th module, and Δ_*k*_ the “imbalance” between the activity of excitatory and inhibitory neurons. In terms of these variables, fixed point equations become

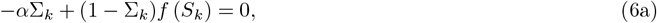

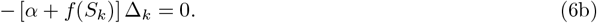

At a fixed point it is therefore Δ_*k*_ = 0, which means that the activity of excitatory and inhibitory neurons inside each module is equal. This is a consequence of the structure of the connections, namely the fact that excitatory and inhibitory neurons of a given module receive the same input. We then make the assumption that

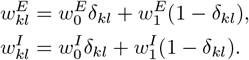

In this case, Eq. (6a) will be given by *F*_*k*_(*{*Σ_*l*_*}*) = 0, where

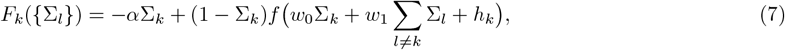

where 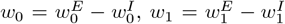. In the following, we consider the external input *h*_*k*_ = 0 for all the modules, and *α* = *β*. Eq. (7) always has the solution with all the activities Σ_*k*_ = 0. This is an absorbing quiescent fixed point, in the sense that the dynamics ceases completely in that state, and has to be restarted by activating one or more neurons by an external input. Depending on the values of *w*_0_ and *w*_1_, this fixed point can be attractive or repulsive. When it is attractive, we call the phase “SL”, which stands for “symmetric low” activity. In this phase, for a very low external input, the activity of the network is correspondingly low and incoherent, with sparse and uncorrelated spikes. For values of *w*_0_ and *w*_1_ for which the quiescent fixed point is repulsive, different attractive fixed points appear, in which the activity is different from zero also in absence of external input, that is the dynamics is self-sustained. We look for fixed points where modules *k* = 1, …, *m* have an activity Σ_*k*_ = Σ *>* 0, while module *k* = *m* + 1, …, *M* have Σ_*k*_ = 0, so that the fixed point equations become, for *h*_*k*_ = 0,

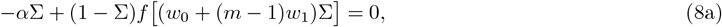

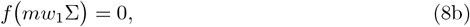

respectively for *k* ≤ *m* and *k > m*. By Eq. (8a), as the function *f* (*s*) is increasing and *f* (*s*) ≤ *βs* for *s >* 0, there is a solution with Σ *>* 0 only for *w*_0_ + (*m* − 1)*w*_1_ *> β*^−1^*α*. On the other hand, if *m < M*, by Eq. (8b) it has to be *w*_1_ ≤ 0. Therefore, for *w*_1_ *>* 0 and *w*_0_ *> β*^−1^*α* − (*M* − 1)*w*_1_, there is only a solution with all the Σ_*k*_ greater than zero. We call this phase “SH”, which stands for “symmetric high” activity. For *w*_1_ ≤ 0 and *w*_0_ ≥ *β*^−1^*α* − (*m* − 1)*w*_1_, there will be also the solution with *m* modules having Σ_*k*_ *>* 0 and *M* − *m* modules having Σ_*k*_ = 0. We call this solution “B*m*”, where the B stands for breaking of symmetry. Of course there are 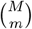 such fixed points, corresponding to which modules are chosen to be active.

To look for the stability of the fixed points, from Eq. (7) one has to compute the Jacobian matrix 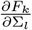, ad its eigenvalues. If all the eigenvalues are negative, then the fixed point is attractive. If on the other hand one or more eigenvalues become positive, then the fixed point becomes repulsive. In Fig. 2 we plot the phase diagram in the case *M* = 3, showing which fixed point is stable depending on the values of *w*_0_ and *w*_1_. The two solid lines, respectively *w*_0_ = *β*^−1^*α* for *w*_1_ ≤ 0, and *w*_0_ = *β*^−1^*α* − (*M* −1)*w*_1_ for *w*_1_ ≥ 0, represent the limit of stability of the low activity fixed point, that is of the phase SL. Exactly on those lines, one of the eigenvalues of the Jacobian matrix becomes equal to zero, so that the stability of the low activity phase is only marginal. If one or more neurons are activated by an external input, then an “avalanche” is generated, that is a very long sequence of activations and deactivations, before the system comes back to the quiescent state. These avalanches have a power law distributions of the durations and sizes, as it will be shown in the next section.

**FIG. 1.**
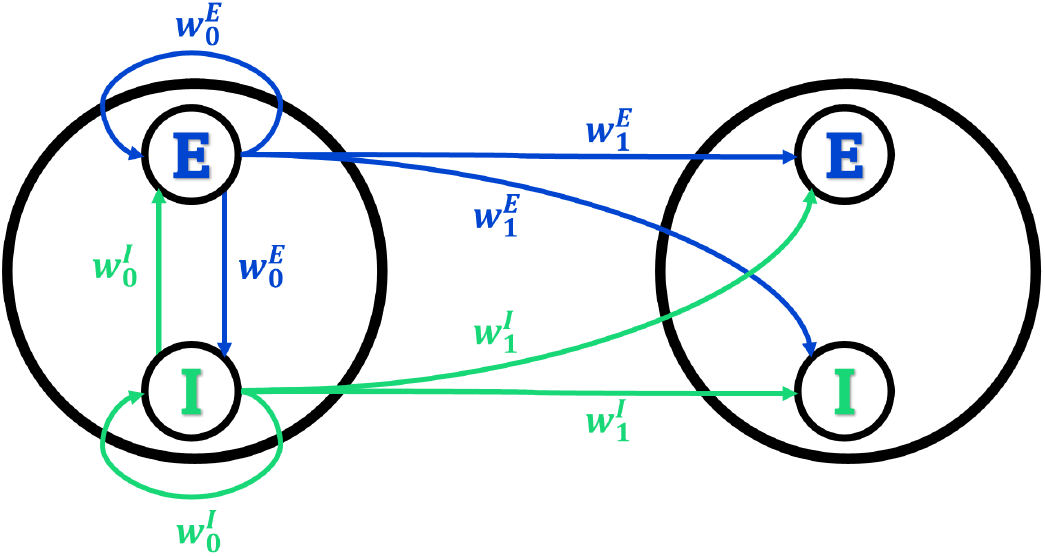
Connectivity of the network. Only the first two modules, and the connections outgoing from neurons of the first module, are shown. The connectivity matrix is symmetric for the permutation of the modules, so that the connections outgoing from any module to any other module are equal.

**FIG. 2.**
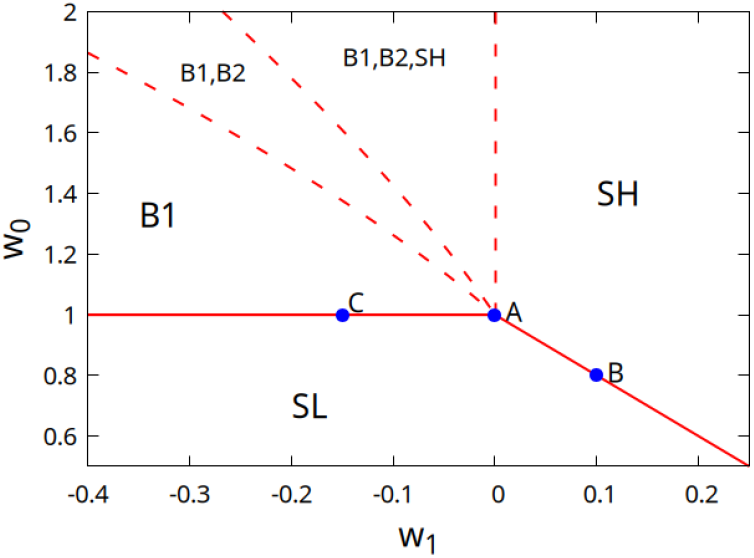
Phase diagram of the Wilson-Cowan model on a modular network, in the case *M* = 3 and for *α* = *β*. In each region of phase space, the phases that are stable (that is whose fixed point is attractive) are listed. In the phase SL the activities of all the modules are low (proportional to *h*) for *h* → 0. In the phase SH the activities of all the modules are high (finite in the limit *h* → 0). In phase B1 (B2) one (two) modules have high activity and the others low activity. The values of the activity are continuous at the boundary of phase SL, as the activity in phases SH and B1 tends to zero when approaching the separation lines, that is transitions are “second order” in the language of statistical physics. The fixed point of the dynamics is “marginally stable” there, so that avalanches have a scale-free distribution. Blue dots (A, B and C) are the points where avalanche distributions and their average shape are investigated in the following section. On the other hand, at dashed lines one or more fixed points lose stability in a discontinuous manner.

## III. CRITICAL DYNAMICS AT THE EDGE OF THE LOW ACTIVITY PHASE

In this section, we study the distribution of bursts of activity (avalanches), and their average shape, varying the strength of the excitatory and inhibitory connections, both between different modules (inter-module connections, labeled by index 1) and within each module (intra-module connections, labeled by index 0). While the fixed points of the dynamics, defined by Eqs. (6), only depend on the “imbalances” 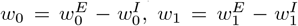, that is the differences between excitatory and inhibitory connections, we will see that the probability distributions of the avalanches and their average shape depend separately on all the parameters 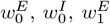 and 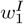. We therefore, for given values of *w*_0_ and *w*_1_, also change the values of the sums 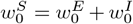 and 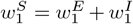. As we have defined 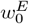, 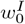, 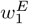 and 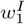 to be non-negative, it has to be 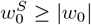 and 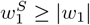.

In the following we consider *M* = 3, *α* = *β* = 0.1 ms^−1^, *N*_*E*_ = *N*_*I*_ = 10^5^. We moreover use *h* = 0. This means that the state where all the neurons are quiescent is an absorbing one, that is the dynamics of the network cease completely. Therefore, each time the network reaches such an absorbing state, we start the dynamics again by setting one neuron of one of the modules (chosen randomly) active “by hand”. The dynamics therefore corresponds to the limit *h* → 0, where after the avalanche is started, neurons can become active only due to the input of the network itself, since there is no external input.

After setting the first neuron active, we simulate the system by means of the event-driven Gillespie algorithm [75]. The steps of the algorithm are the following: 1) for each neuron *i* compute the transition rate *r*_*i*_: *r*_*i*_ = *α* if neuron *i* is active, or *r*_*i*_ = *f* (*s*_*i*_) if it is quiescent; 2) compute the sum over all neurons *r* = ∑_*i*_ *r*_*i*_; 3) draw a time interval *dt* from an exponential distribution with rate *r*; 4) choose the *i* −th neuron with probability *r*_*i*_*/r* and change its state; 5) update the time to *t* + *dt*. The avalanche ends when all the neurons become quiescent. The size of the avalanche is defined as the total number of spikes, that is transitions of a neuron from the quiescent to the active state. The duration is the difference in time between the first spike (the first neuron that is activated “by hand”), and the last deactivation of a neuron.

### A. Disconnected modules

In this subsection, we start by considering the case *w*_1_ = 0, 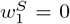, that corresponds to 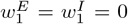. This is therefore the case of disconnected modules, that coincides with the case of a single module *M* = 1, that is with a homogeneous (non-modular) network, already studied in [29]. In this case we have just two phases (two attractive fixed points), corresponding to a phase of low incoherent activity for *w*_0_ *< w*_0*c*_, and a phase of high self-sustained activity for *w*_0_ *> w*_0*c*_, where 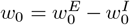 is the “imbalance” between excitatory and inhibitory connections. These phases correspond to the phases SL and SH of the modular network. The critical point is given by *w*_0*c*_ = *β*^−1^*α*, that is *w*_0*c*_ = 1 as we consider here *α* = *β*. For *w*_0_ *< w*_0*c*_, the attractive fixed point corresponds to a quiescent network, while for *w*_0_ *> w*_0*c*_ the attractive fixed point corresponds to a finite value of the activity, so that the dynamics of the network is self-sustained. At *w*_0_ = *w*_0*c*_ = 1 the quiescent state is marginally stable, which means that an avalanche started by setting just one neuron active will have a power law distribution of sizes and durations, with a cut-off only determined by the finite number of neurons. The point *w*_0_ = 1, *w*_1_ = 0 corresponds to the point “A” in Fig. 2. Note that in the case of disconnected modules (*w*_1_ = 0, 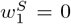), and in general in the case *w*_1_ ≥ 0 (balanced or “excitatory imbalanced” inter-module connections), there can be no “broken symmetry” phases B*m*, that appear only for “inhibitory imbalanced” (*w*_1_ *<* 0) inter-module connections.

In Fig. 3 we show the avalanche statistics for different values of 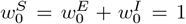, 2, 5, 10. The case 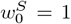 corresponds to a value 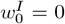 of the inhibitory connections, that is to a purely excitatory network. In Fig. 3A and B we show the probability distribution of avalanche sizes and durations, while in Fig. 3C we show the average size of the avalanche as a function of its duration. Dashed lines correspond to the predictions of the branching model, or mean-field directed percolation (MFDP). It is interesting to note that the larger the value of 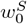, the smaller the deviations with respect to the branching model predictions. The purely excitatory network, while converging to the predicted exponents for very large avalanches, has the most significant deviations for not very large ones. In Fig. 3D we show the mean skewness of the avalanches as a function of their duration. The skewness is strictly zero for the purely excitatory network, and increases for durations around *T* = 100 ms, when increasing the strength of both inhibitory and excitatory connections. (Note that the difference 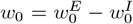 has to be kept constant, in order to remain in the critical point.) Finally in Fig. 3E and F we show the average shape of the avalanches, that is the average number of spikes per millisecond as a function of the time since the start of the avalanche, for avalanches of duration *T* = 100 and *T* = 1000 ms. Notably, the single module only produces symmetric avalanches (in the purely excitatory case), or avalanches with a rightward asymmetry for an intermediate range of durations, with the skewness increasing with the increase of the strength of the connections (and thus of the relative strength of inhibitory connections).

**FIG. 3.**
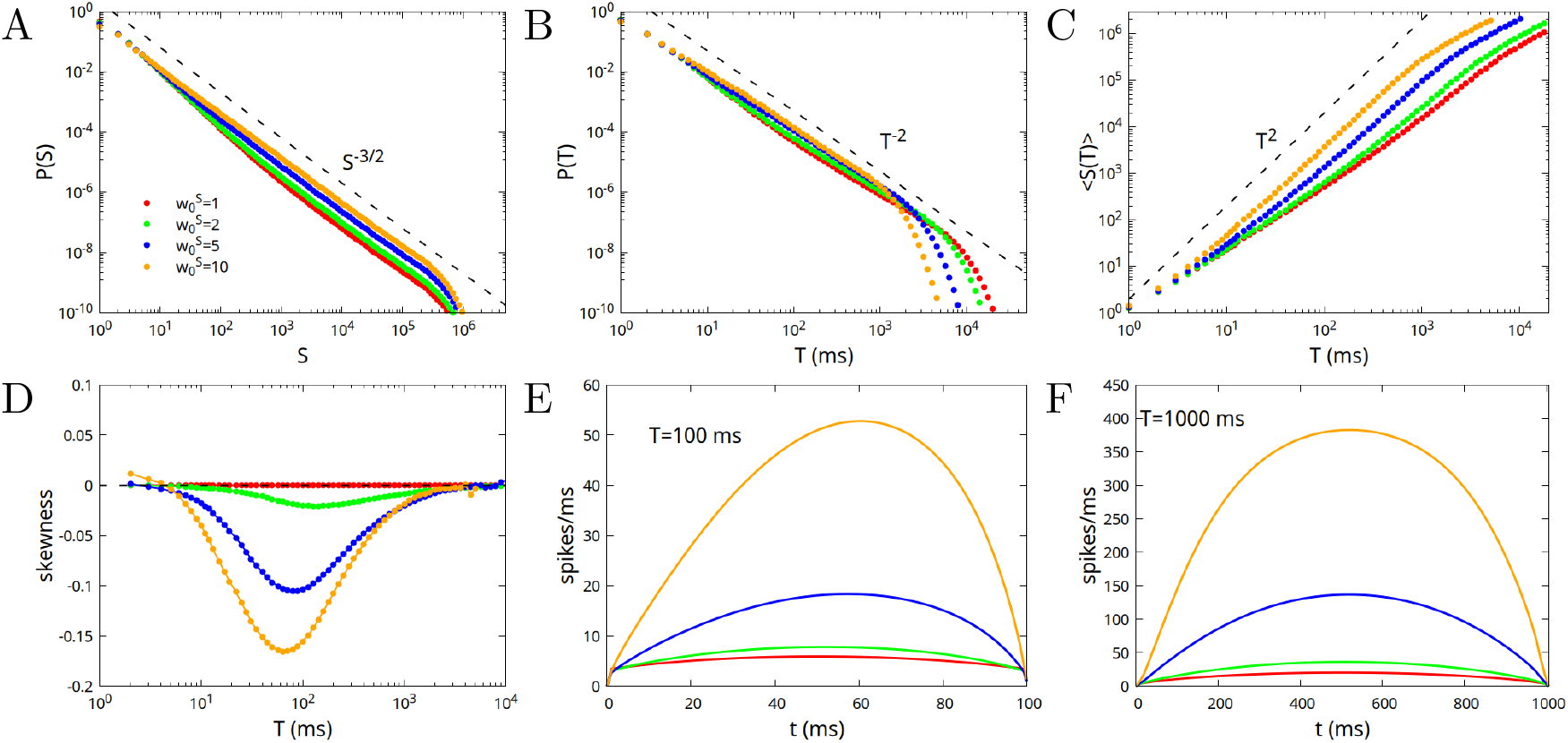
Avalanche statistics in the case of disconnected modules, or of a single module, for 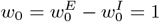(critical value), and different values of the parameter 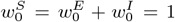, 2, 5, 10. The case 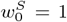 corresponds to a value 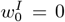 of the inhibitory connections (purely excitatory network), and is characterized by a perfect symmetry of the avalanches, with skewness equal to zero. A) Probability distribution of the sizes; B) Probability distribution of the durations; C) Mean size as a function of the duration; D) Mean skewness of the avalanches as a function of the duration; E) Average shape of the avalanches as a function of time, for avalanches of duration *T* = 100 ms; F) Same as E but for durations *T* = 1000 ms.

In conclusion, in a range of intermediate durations 10 *< T <* 1000, for small values of 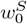 we find nearly symmetric avalanches but strong deviations from MFDP exponents in the distribution of avalanches. On the other hand, for high values of 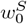, we find very good agreement with MFDP exponents, but right-skewed avalanches. This is quite counterintuitive, as avalanches in MFDP have a symmetric (inverted parabolic) shape.

### B. Modular network with balanced inter-module connections

In this subsection, we consider the case in which the system is at the point *w*_0_ = 1, *w*_1_ = 0 (as in the previous subsection), corresponding to the crossing of all the lines in Fig. 2 (point “A”), on the boundary between phase SL and the different phases of high activity, but this time 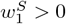. The modules that constitute the network are therefore connected by excitatory and inhibitory connections greater than zero but exactly balanced, that is with 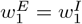. We fix the value 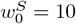, and consider different values of 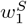. As previously remarked, the fixed points of the dynamics, and their stability, depend only on the “imbalances” 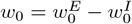 and 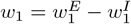, so that once *w*_0_ and *w*_1_ are fixed to the critical point, changing the values of 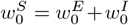 and 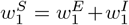 cannot bring the system away from criticality. Moreover, it can be expected that, for a very large system and for very long avalanches, the exponents of power law distributions, and their average shape, do not depend on the values of 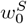 and 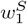. However, their value can change the probability distributions and the shape of not very large avalanches. This was already observed in Fig. 3 for a disconnected network (or a single module) with 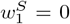, and is confirmed in Fig. 4 for a connected network formed by *M* = 3 modules and 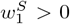. Deviations from MFDP behaviour are largest for small values of 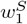 (small values of both excitatory and inhibitory inter-module connections), and tend to reduce for larger values of 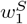. Moreover, while very large avalanches tend to be symmetric, intermediate ones with durations 10 *< T <* 1000 are generally right-skewed (negative skewness), apart from the case 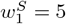 that shows avalanches slightly left-skewed for durations around *T* = 1000.

**FIG. 4.**
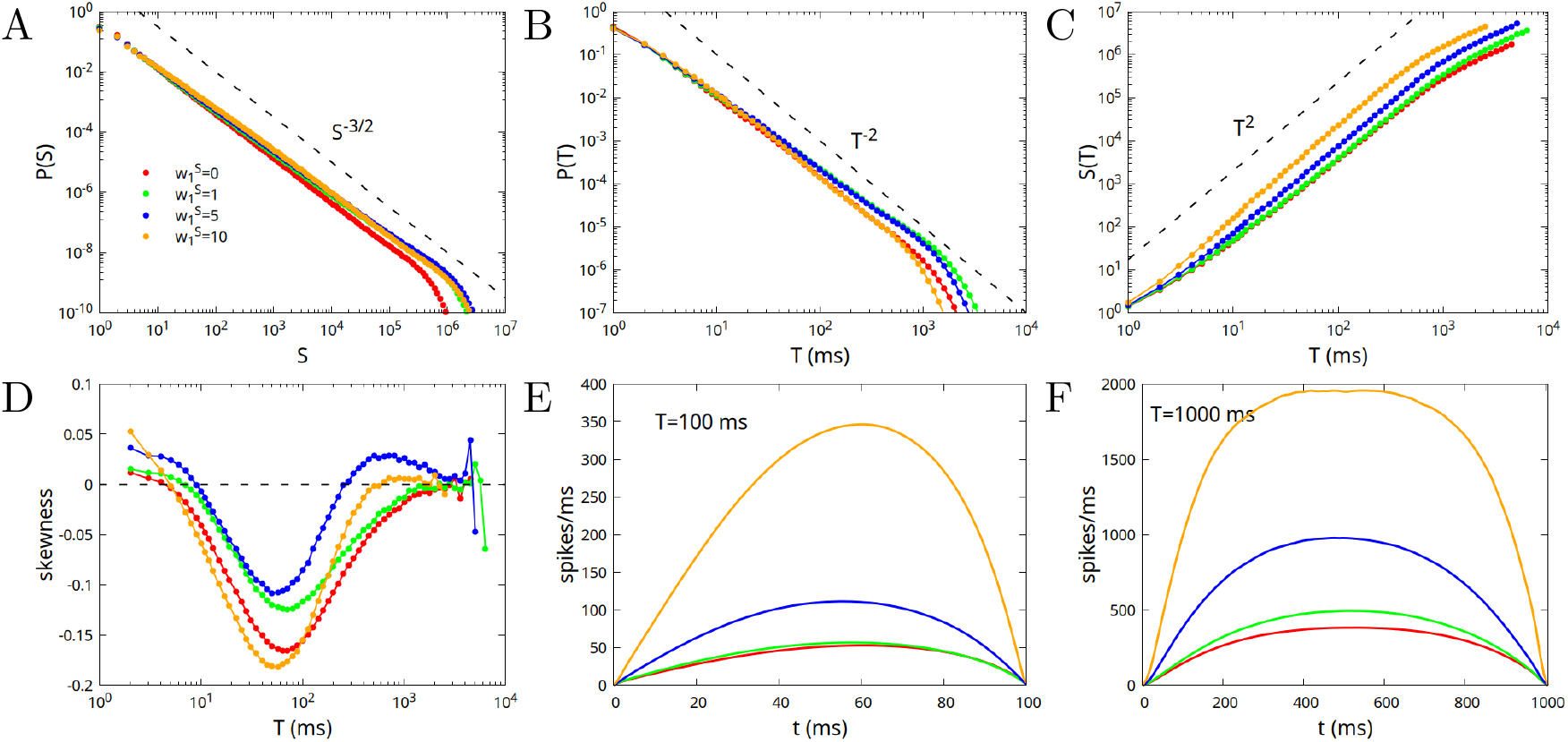
Avalanche statistics in the case of a network constituted by *M* = 3 modules, at the critical point *w*_0_ = 1, *w*_1_ = 0, so that inter-module excitatory and inhibitory connections are exactly balanced, 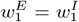. The value of 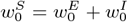 is fixed to 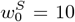, while 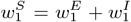 varies between 1 and 10. For reference, we include in the plots also the case 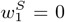 (red lines), that corresponds to the yellow lines in Fig. 3. A) Probability distribution of the sizes; B) Probability distribution of the durations; C) Mean size as a function of the duration; D) Mean skewness of the avalanches as a function of the duration; E) Average shape of avalanches as a function of time, for avalanches of duration *T* = 100 ms; F) Same as E but for durations *T* = 1000 ms.

### C. Modular network with “excitatory imbalanced” inter-module connections

If we shift the balance of inter-module connections making them excitatory imbalanced, that is take *w*_1_ *>* 0, we have to decrease correspondingly the imbalance of intra-module connections to remain at the critical point. Indeed as discussed after Eqs. (8) and shown in Fig. 2, the critical line for *w*_1_ ≥ 0 is given by the equation

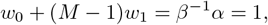

having chosen *α* = *β*. In Fig. 5 we show the case *M* = 3, *w*_1_ = 0.1, *w*_0_ = 0.8, corresponding to the point “B” in Fig. 2. As can be seen by comparing Figs. 4 and 5, the behaviour for *w*_1_ *>* 0 is quite similar to that for *w*_1_ = 0. There are deviations from the MFDP predictions at low values of 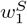, that are reduced increasing 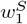. Moreover, for large values of 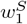, avalanches become right-skewed for intermediate values of the duration.

**FIG. 5.**
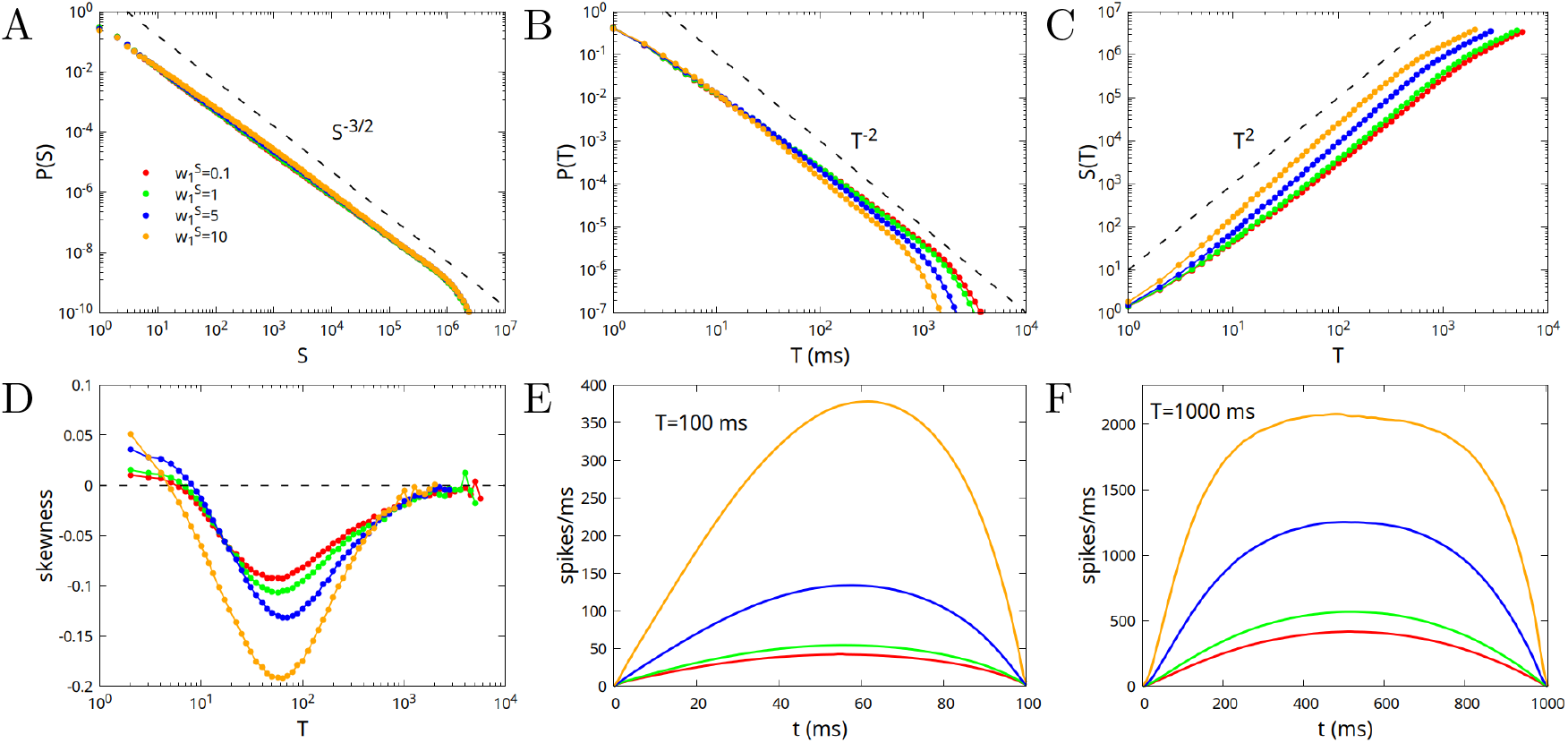
Avalanche statistics in the case of a network constituted by *M* = 3 modules, at the critical point *w*_0_ = 0.8, *w*_1_ = 0.1 (point “B” in Fig. 2), that is for “excitatory imbalanced” inter-module connections. The value of 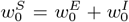 is fixed to 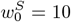, while 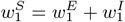 varies between 0.1 and 10. A) Probability distribution of the sizes; B) Probability distribution of the durations; C) Mean size as a function of the duration; D) Mean skewness of the avalanches as a function of the duration; E) Average shape of avalanches as a function of time, for avalanches of duration *T* = 100 ms; F) Same as E but for durations *T* = 1000 ms.

### D. Modular network with “inhibitory imbalanced” inter-module connections

As shown in Fig. 2, if inter-module connections are “inhibitory imbalanced”, that is 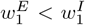 and *w*_1_ *<* 0, then “broken symmetry” phases appear, where a subset of the modules have high activity, while others have low activity. In particular, symmetric phase SL becomes unstable at the critical line *w*_0_ = 1 toward phase B1, where only one of the modules is active. In Fig. 6 we show avalanche statistics for *w*_0_ = 1, *w*_1_ = −0.15, that is at point “C” in Fig. 2, for 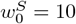 and different values of 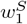 between 0.5 and 10.

**FIG. 6.**
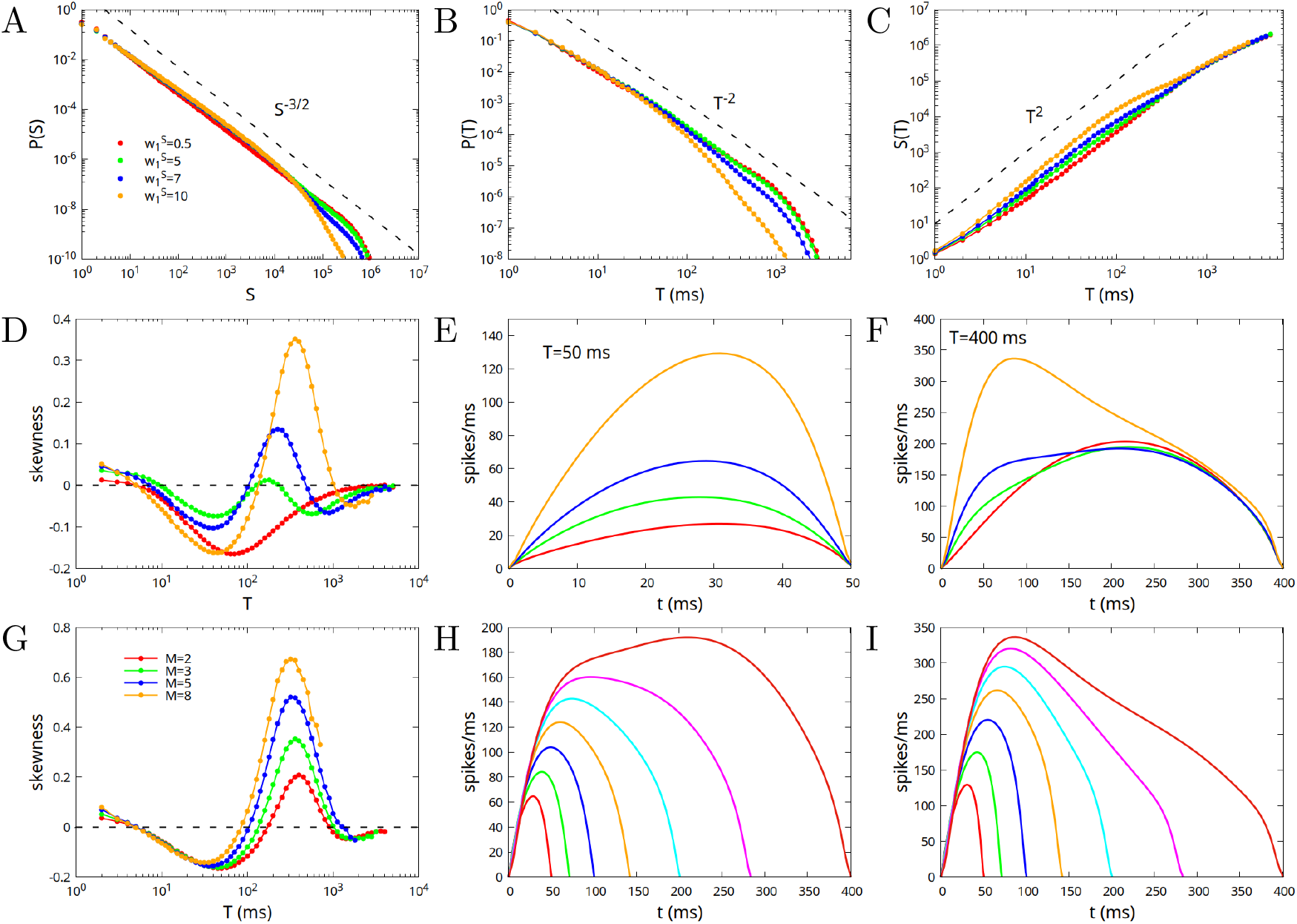
Avalanche statistics in the case of a network constituted by *M* = 3 modules, at the critical point *w*_0_ = 1, *w*_1_ = 0.15 (point “C” in Fig. 2), that is for “inhibitory imbalanced” inter-module connections. The value of 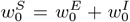 is fixed to 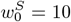, while 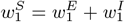 varies between 0.5 and 10. A) Probability distribution of the sizes; B) Probability distribution of the durations; C) Mean size as a function of the duration; D) Mean skewness of the avalanches as a function of the duration; E) Average shape of avalanches as a function of time, for avalanches of duration *T* = 50 ms; F) Same as E but for durations *T* = 400 ms; G) Average shape of avalanches of different durations, for 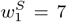 H) Same as G but for 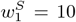 I) Mean skewness of the avalanches as a function of the duration, for different number of modules, 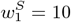.

As shown in Fig. 6A-C, in this case, differently from the case *w*_1_ *>* 0, the largest deviations from MFDP are observed at large values of 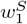, in particular between *T* = 100 ms and *T* = 1000 ms, where both the distributions of sizes *P* (*S*) and of durations *P* (*T*) decay with an exponent larger (steeper decay) than the one of MFDP, while the average size as a function of duration ⟨*S*⟩ (*T*) increases with a smaller exponent. Notably, in the same range of avalanche durations, that is between *T* = 100 ms and *T* = 1000 ms, the average shape of the avalanches tends to become leftward skewed (positive skewness) as the strength of inter-module interactions increases, that is for larger values of 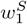, as shown in Fig. 6D. Figs. 6E and F show the average shape for avalanches of duration 50 ms and 400 ms. For 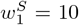 and durations around 400 ms, the skewness assumes a quite large value of almost 0.4. In Fig. 6G and H we plot the average shape for different avalanche durations, respectively for 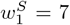 and 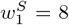. Finally in Fig. 6I we plot the skewness as a function of the number of modules, showing that the effect is enhanced for greater number of modules. In the followinf subsection, we investigate the origin of such a leftward asymmetry, given that similar results have been reported in several experimental systems, as discussed in the Introduction. Specifically, we analyze more in detail the differences between avalanches for “excitatory imbalanced” and “inhibitory imbalanced” inter-module connections, that is at points B and C of Fig. 2.

### E. Comparison between “excitatory imbalanced” and “inhibitory imbalanced” inter-module connections

In Fig. 7 we show different examples of avalanches of duration *T* = 400 ms, in the case of “excitatory imbalanced” (upper row, parameters *w*_0_ = 0.8, *w*_1_ = 0.1) and “inhibitory imbalanced” (lower row, parameters *w*_0_ = 1, *w*_1_ = −0.15) inter-module connections, highlighting the contributions of the individual modules to the avalanche. The most apparent feature of the plots is the fact that, in the first case where inter-module connections are “excitatory imbalanced”, there is a strong positive correlation between the profiles of the avalanche across all three modules. It is quite predictable then that the profile is not very different from the one observed in the case of a single module, or disconnected modules, that is with a rightward asymmetry. On the other hand, in the case of “inhibitory imbalanced” inter-module connections, the behaviour is markedly different. We observe a first interval of time, of about 100 ms, where the avalanche spreads in a nearly uniform way in the three modules, while for subsequent times one of the modules prevails over the others, and remains almost the only active module until the end of the avalanche. To understand the origin of this behaviour, we plot in Fig. 8A the average imbalance ⟨Δ_*k*_⟩ = ⟨ (*x*_*k*_ −*y*_*k*_)*/*2⟩ between the excitatory activity *x*_*k*_ (fraction of active excitatory neurons) and the inhibitory activity *y*_*k*_, on the *k*-th module, and the average total activity ⟨Σ_*k*_⟩ = ⟨ (*x*_*k*_ +*y*_*k*_)*/*2⟩ (Inset of Fig. 8A). As ⟨· · · ⟩ represents the average over all the avalanche of fixed duration, ⟨Δ_*k*_⟩ and ⟨Σ_*k*_⟩ are equal for all the modules *k* = 1, … *M*. Red (orange) line corresponds respectively to the excitatory (inhibitory) imbalanced case, that is to parameters *w*_0_ = 0.8, *w*_1_ = 0.1 (*w*_0_ = 1, *w*_1_ = −0.15). From these, one can compute the “intra-module” and “inter-module” input of neurons. The total input *s*_*k*_ of a neuron in module *k*, given by Eq. 3, can indeed be written as *s*_*k*_ = *s*_0,*k*_ + *s*_1,*k*_ + *h*_*k*_, where

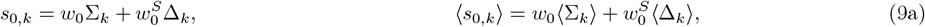

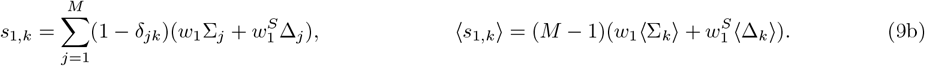

**FIG. 7.**
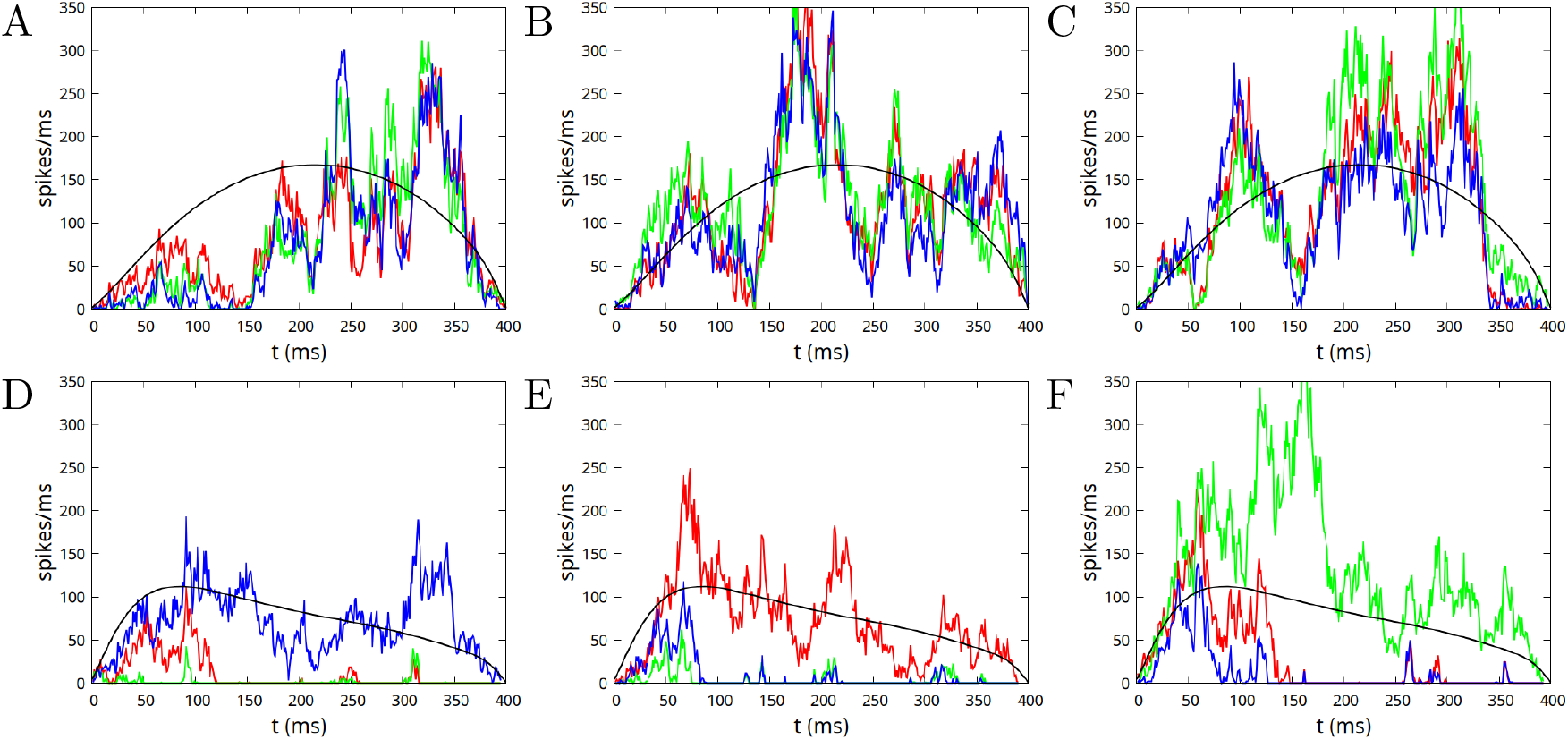
Examples of avalanches of duration *T* = 400 ms, in the case of “excitatory imbalanced” (upper row, parameters *w*_0_ = 0.8, *w*_1_ = 0.1) and “inhibitory imbalanced” (lower row, parameters *w*_0_ = 1, *w*_1_ = −0.15) inter-module connections. In both cases 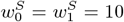. The istantaneous firing rate for three different realizations of an avalanche (on the three columns) are shown for the *M* = 3 modules with respectively red, green and blue lines. Black lines show the average firing rate (over all avalanches with duration 400 ms), on a single module in the first case (upper row), and the total average rate on the three modules in the second case (lower row).

**FIG. 8.**
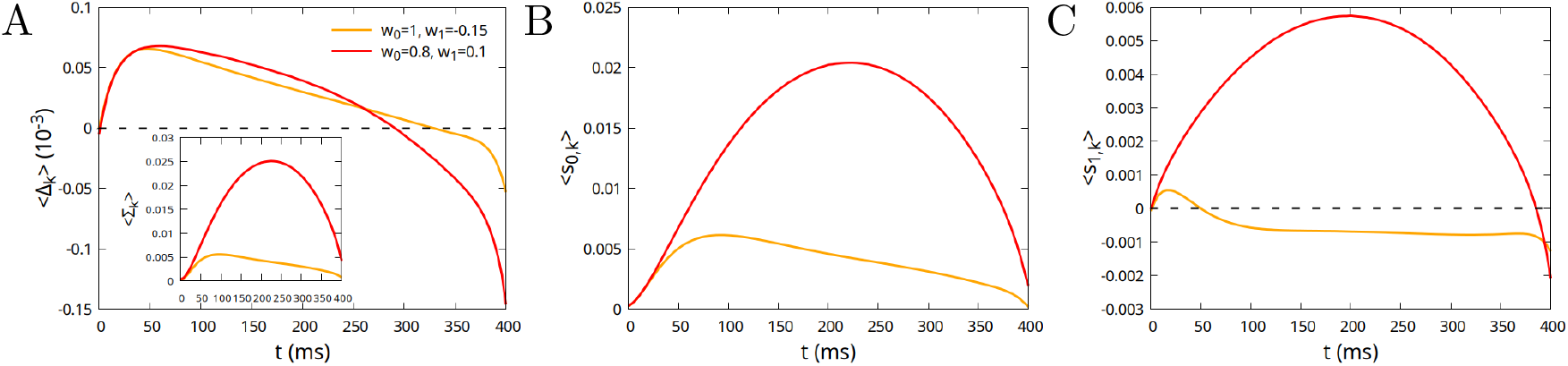
(A) Average half-difference ⟨Δ_*k*_⟩ between the excitatory and inhibitory activity (fraction of active neurons) on the k-th module during an avalanche of duration *T* = 400 ms, in the case of “excitatory imbalanced” (red line) and “inhibitory imbalanced” (orange line) inter-module connections. Inset: average half-sum ⟨Σ_*k*_⟩ of the activities; (B) Average input to the neurons from the same module to which they belong; (C) Average input to the neurons from the modules to which they do not belong.

The input *s*_0,*k*_ comes from neurons belonging to the same module, while *s*_1,*k*_ from neurons belonging to different modules. In the case *w*_1_ *<* 0 (“inhibitory imbalanced” inter-module connections) the input coming from module *k* to neurons in different modules can still be positive, if

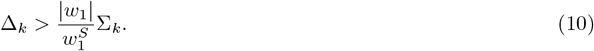

The two contributions ⟨*s*_0,*k*_⟩ and ⟨*s*_1,*k*_⟩ are plotted in Fig. 8B and C. It can be noticed that, in the case of “inhibitory imbalanced” inter-module connections (orange lines), the external input (Fig. 8C) is positive in a first interval of around 50 ms, due to the prevalence of the activity of excitatory neurons in the beginning of the avalanche, so that Eq. (10) is satisfied. Therefore, although inter-module inhibitory connections are stronger than inter-module excitatory ones, in the beginning of the avalanche one observes a “mutual reinforcement” between the modules, as the one observed in the case of excitatory imbalanced connections (albeit of course weaker). Only for times larger than 100–150 ms mutual inhibition, and therefore the symmetry breaking between modules, takes hold. This explains why leftward avalanches are observed only for durations between 100 and 1000 ms. For shorter durations, symmetry breaking has not developed yet. On the other hand, for durations larger than 1000 ms, the initial time window where all the modules are engaged in the avalanche can be neglected. For intermediate durations, the avalanche is formed by a first interval where all the modules are engaged, and a second one where only one module is left, giving rise therefore to a leftward asymmetry.

## IV. CONCLUSIONS

Modularity, likely shaped by evolutionary and functional constraints, represents a fundamental organizing principle of complex networks across social, technological, and biological systems [76, 77]. This principle is observed across levels of biological organization, from genetic regulatory and protein interaction networks to the structure of ecosystems. A modular community architecture constitutes a core feature of both functional and structural brain networks [62, 63, 78, 79], observed at both whole-brain and cellular scales. Here, we have studied a stochastic Wilson-Cowan model incorporating a modular architecture, to investigate how such structural organization influences spontaneous dynamics, exploring the phase diagram and the avalanche shape near the analytically derived critical lines.

Experimentally, avalanche shapes can exhibit marked left-skewed asymmetry, that are not reproduced in stochastic Wilson-Cowan models lacking modular network organization, which, both under all-to-all coupling and within two-dimensional network structures, display either symmetric or rightward avalanche shapes in the vicinity of the critical point [29, 30].

To clarify the role of network architecture, while capturing the modular organization observed in the brain, we investigated a stochastic modular Wilson-Cowan model composed of *M* modules, coupled by intra and inter-module connections, under a vanishing external drive. By analyzing the fixed-point equations and linearizing the dynamics around them, we derived a phase diagram comprising symmetric phases with low (SL) or high (SH) activity in all modules, and broken symmetry phases B*m*, where only *m < M* modules are active. Along two critical lines, between phases SL and SH, and between SL and B1, the distribution of the avalanches is scale-free.

We then characterized the average shape of the avalanches along these lines. At the critical line between phases SL and SH, avalanches show a rightward shape, as in the case of a non-modular architecture. On the other hand, at the critical line between SL and B1, that is at the edge of the broken symmetry phase, avalanches show a leftward asymmetry for an intermediate range of durations. We have analyzed in detail the dynamics of the network during an avalanche in this latter condition, finding that it proceeds in two stages: an initially cooperative regime, where excitatory activity is prevalent, followed by inhibitory competition that selects one dominant module and suppresses the others. This is the relevant mechanism leading to a fast rise of the avalanche, followed by a slower decay, and therefore to leftward asymmetry.

The role of inhibition in producing leftward-skewed shapes has also been recently found [80] in an integrate-and-fire model with short- and long-term plasticity and more highly connected inhibitory neurons, which highlighted the crucial role of inhibition, and of differences in synaptic recovery rates between excitatory and inhibitory neurons, in generating leftward asymmetry.

In conclusion, this model demonstrates that the leftward shape of avalanches emerges as a direct consequence of the symmetry-breaking transition driven by competitive inter-module connections, where inhibition dominates over excitation between modules. Notably, this framework enables systematic manipulation of inter-module interactions, biasing them toward either competitive (inhibitory) or cooperative (excitatory) dynamics, while maintaining critical behavior by adjusting the E/I balance of intra-module connections. Together, these findings highlight how the interplay between intra- and inter-module connectivity shapes both the nature of the symmetry-breaking transition and the emergence of asymmetric avalanches, providing a mechanistic link between network architecture and spontaneous neural dynamics.

## Notes

### Competing Interest Statement

The authors have declared no competing interest.

